## Supplementary Information for "Intermolecular Channels Direct Crystal Orientation in Mineralized Collagen"

**This PDF file includes:**

Supplementary Information 1 to 14
Captions for Movies S1 to S8
References for SI reference citations

**Other supplementary materials for this manuscript include the following:**

Movies S1 to S8

#### Supporting Information 1

**Overview of the human bone lamella**

Figure S1 is a 2D projection of tomographic reconstruction slices (70 z-slices averaged), showing an overview of the human bone lamella. Most of the area shows a clear ~67 nm banding pattern, indicating that collagen fibrils are closely packed, and their long axes are generally in plane. Gap regions are darker than the overlap regions due to heavy metal staining of the sample (using the OTOTO method, similar to Landis using uranyl acetate[1,2]), not due to mineralization, as no higher mineral density is observed. Indeed, the *in vitro* HAp/collagen (Figure 2i), which was only lightly stained, does not show this an effect. In some areas (highlighted by yellow arrows) the banding pattern is less clear, indicating that the long axes of collagen fibrils within these areas are tilted out of plane, as a result of the 3D twisting of collagen fibrils. The area in the yellow box is selected for analysis in Figure 2 in the main text, as a clear discontinuity can be seen between the two adjacent collagen fibrils within this area, which helps to identify the boundary between the two fibrils.

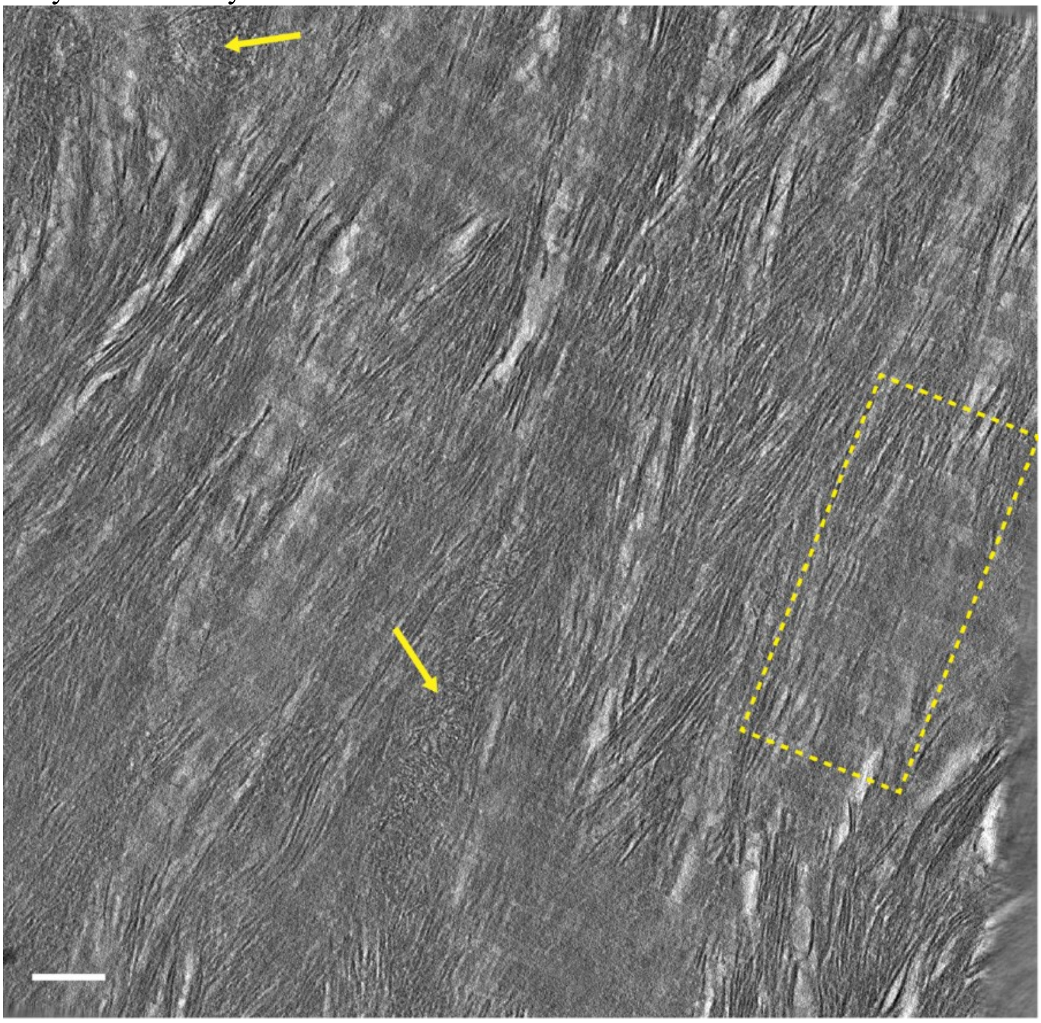

**Figure S1.** 2D projection of tomographic reconstruction slices (70 z-slices averaged), showing an overview of the human bone lamella. Scale bar: 100 nm.

### Supporting Information 2

**Determination of crystal orientations and dimensions inside collagen**

To determine the dimensions of the crystallites that were grown within the collagen a size analysis was performed. To that end tomographic slices were loaded as images into Matlab and the long and short axis were manually determined by clicking the four edges of the crystallites. The slices from the tomogram were taken such that the crystallites were at least 10 nm (in y-direction) or 50 nm (in z-direction) apart so the same crystals are not measured more than once. The contrast of the z-slices was enhanced by averaging 10 adjacent slices. A sample of the size analysis that was performed on different sections for both the mineralized collagen fibril from human bone and the *in vitro* mineralized collagen fibril is shown in Figure S2. Dark streaks are visible in the z-slices (as pointed out by yellow arrows in Figure S2b and d) due the limited tilt increment (2° per step) and tilt range (−65 to 65°) used for collecting the tomography tilt series.[3] These streaks are however low in contrast and each of them only exist within several slices (see Supporting Movies S3 and S5), while a HAp crystal will propagate through ~85 slices due its length (~65 nm). This allows to confidently distinguish between the streaks and HAp crystals.

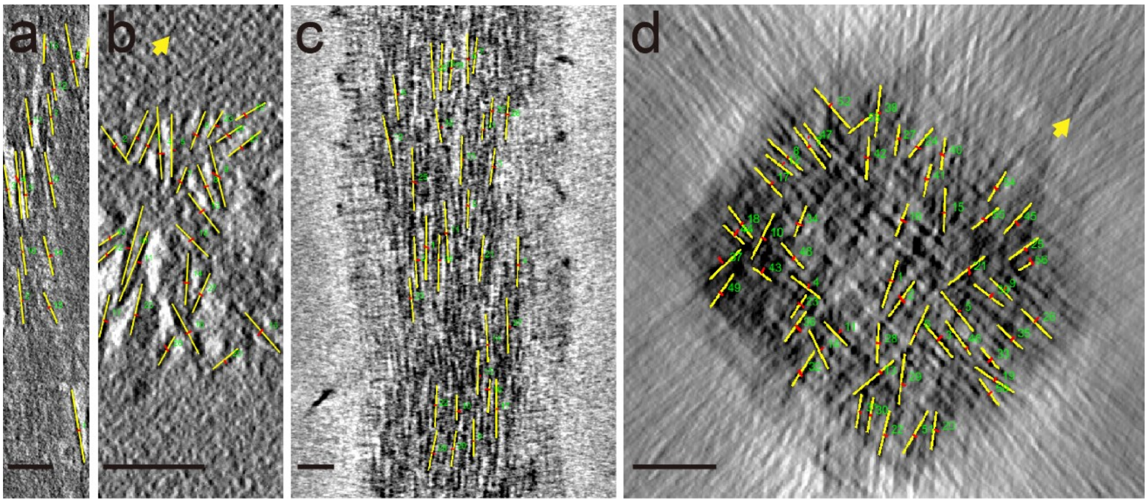


**Figure S2.** Size analysis on the tomographic reconstruction slices, in which the length of long (yellow lines) and short axes (red lines) of the crystals are determined. (a,b) Mineralized collagen fibril in human bone viewed from the top and along the fibril, respectively (c,d) collagen fibril mineralized *in vitro* by HAp viewed from the top and along the fibril, respectively. Dark streaks are artifacts induced by limitation of tilt conditions and are highlighted by yellow arrows in b) and d). Scale bars: 50 nm

#### Supporting Information 3

##### HAp platelets near the collagen fibril boundary

Figure S3 tracks one HAP platelet through different cross section slices of two adjacent collagen fibrils (as shown in Figs. 2a and 2e). The distance between Figures S3b (slice 183) and S3e (slice 253) is 52.5 nm (70 slices, slice thickness=0.75nm). The platelet extends from the boundary between the two fibrils into the fibril on the right side. Note that the lateral orientation of the platelet has rotated counter clockwise with ~30$^{\circ}$ between Figures S3b and S3e, indicating that the platelet is twisted like a propeller as previously reported.[4] Such twisting can also be observed for mineral platelets that are completely within the fibrils.

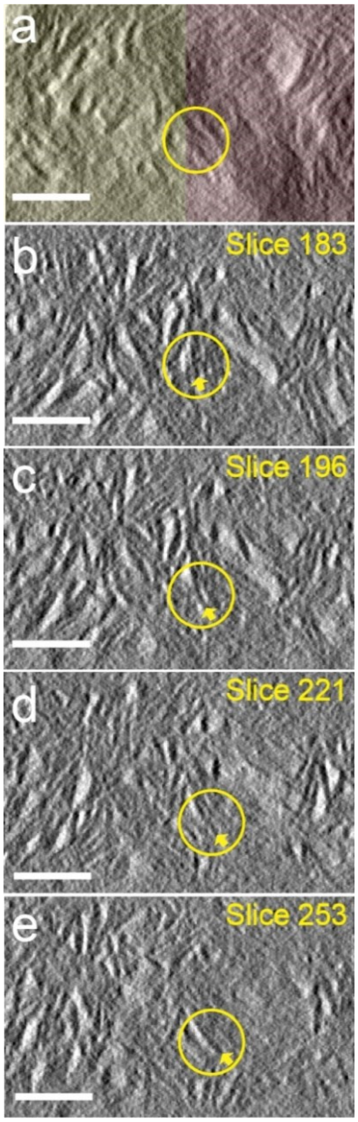


**Figure S3.** (a) 2D projection of 400 cross section tomographic reconstruction slices (total thickness 300 nm) showing two adjacent fibrils, as highlighted by yellow and red colors, respectively. (b-e) Cross section slices of the two fibrils with slice number 183, 196, 221 and 253, respectively. Each slice is averaged from 10 adjacent slices to enhance the contrast. The cross sections of one same HAp platelet in the four slices are identified by yellow circles, while the lateral orientation of the platelet is highlighted by yellow arrows. The position of yellow circle in (b) is also highlighted in (a). Scale bars: 50 nm.

#### Supporting Information 4

**Detailed analysis of the cross section tomography reconstruction slices**

Figure S4 shows 6 cross section tomography reconstruction slices selected at different regions of the two adjacent fibrils. Randomly oriented stacks of 2-4 HAp platelets can be seen in most of the slices, while in the last slice (Figure S4g) there are several larger stacks of ~8 platelets. These stacks are ~50 nm in size and have different lateral orientations. The dots highlighted by yellow circles correspond to the needles found at the tip of platelets.

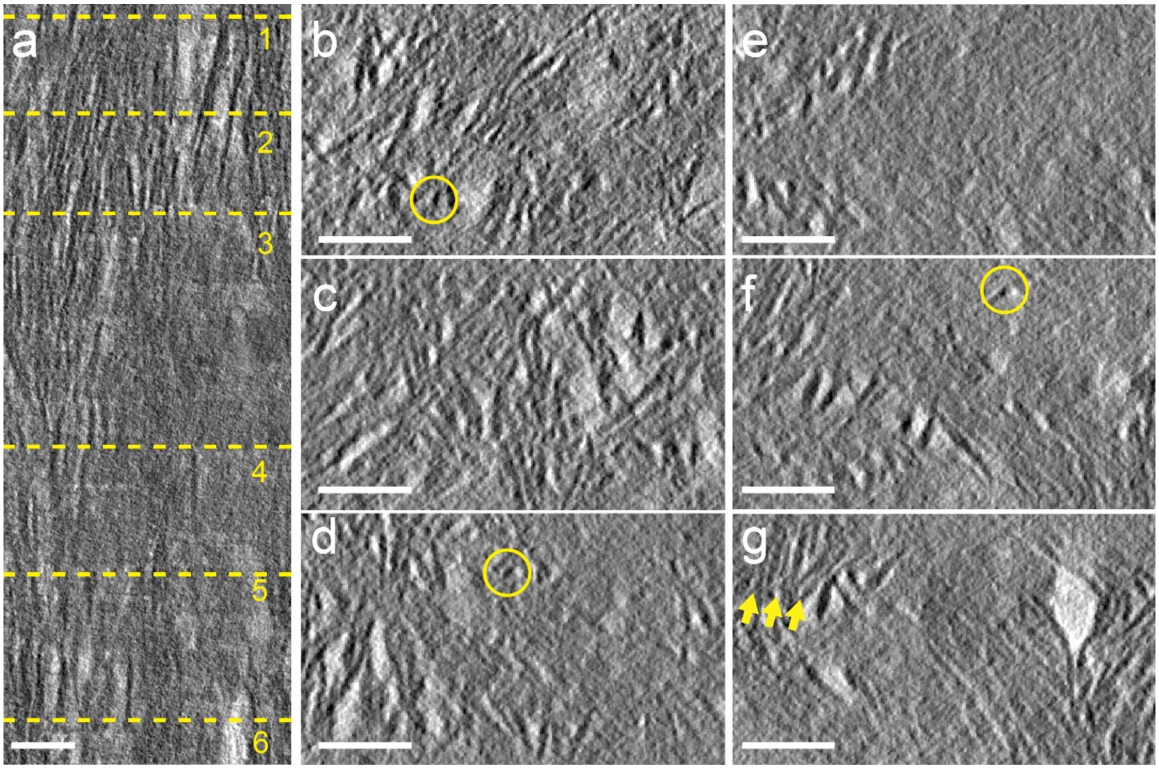


**Figure S4.** (a) 2D projection of the top-view tomographic reconstruction slices showing two adjacent mineralized collagen fibrils in human bone. (b-g) Cross section reconstruction slices at positions 1-6, respectively, as labelled in (a). Each slice is averaged from 10 adjacent slices to enhance the contrast. Several dots correspond to the needle-shaped tips of the mineral platelets are highlighted by yellow circles, while several stacks consist of ~8 platelets are highlight in (g) by array of yellow arrows. Scale bars: 50 nm.

#### Supporting Information 5

**Needle-shaped tips of the HAp platelets**

By tracking a 'dot' in different cross section slices (Figure S5), it is clear that the dot actually corresponds to a needle which is ~2 nm in width. The needle evolves into a platelet-like shape in Figure S5d, suggesting that its length is about ~10 nm (spanning 14 slices, slice thickness 0.75 nm), and the needle is actually the tip of a HAp platelet.

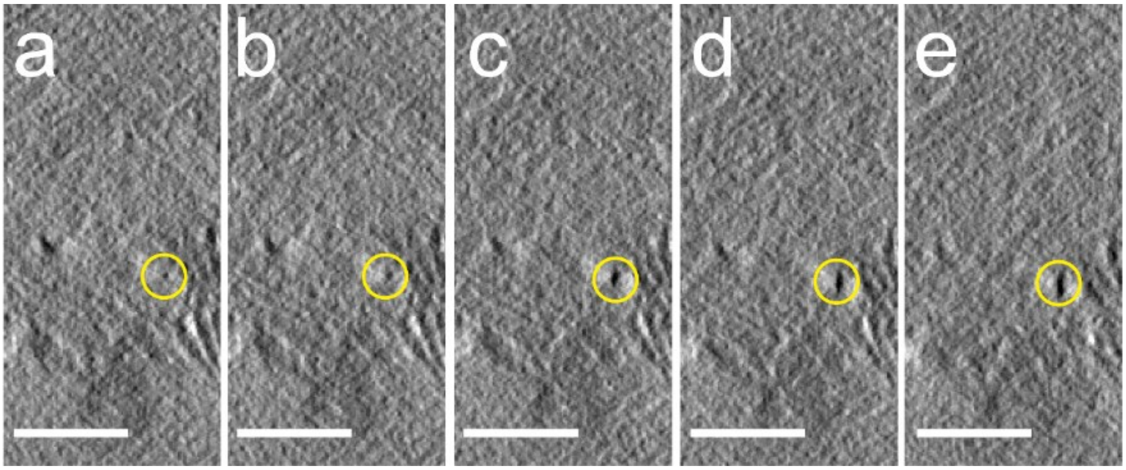


**Figure S5.** (a-e) Cross section slices of one collagen fibril with slice number 579, 574, 568, 566 and 553, respectively. Each slice is averaged from 10 adjacent slices to enhance the contrast. The cross-sections of the needle which evolves into platelet are highlighted by yellow circles. Scale bars: 50 nm.

#### Supporting Information 6

##### Comparison with previously reported tomography results

As our results contrast the early results of Landis et al. on turkey tendon[2] and chick bone,[1] we discuss here the origin of these differences, and propose that they originate from a difference in sample preparation and visualization. As shown in Figure S6, the tomography samples consist of mineralized collagen fibrils aligned within a ~100 nm slab of epoxy-embedded bone material. At low tilting angle, e.g., 0°, the platelets with edge-on orientation are high in contrast due to their widths (18.3 ± 4.3 nm as we observed), while the platelets in other orientations ("*non-edge-on"* platelets) are less visible as they are very thin (2~4 nm). At higher tilting angles, e. g., 60°, the projected sample thickness $T_2$ is doubled with respect to the projected thickness at 0° ($T_1$). This reduces the contrast of all platelets with respect to the matrix at higher tilt angles, even more so when the sample is relatively thick. Therefore, the "*non-edge-on"* platelets that would to be more visible at these higher tilt angles are still low in contrast. As a result, the parallel organized platelets that are edge-on at low tilting angles will be significantly more visible in the final reconstruction than the "*non-edge-on"* platelets, thereby generating the impression of a "deck of cards" organization.

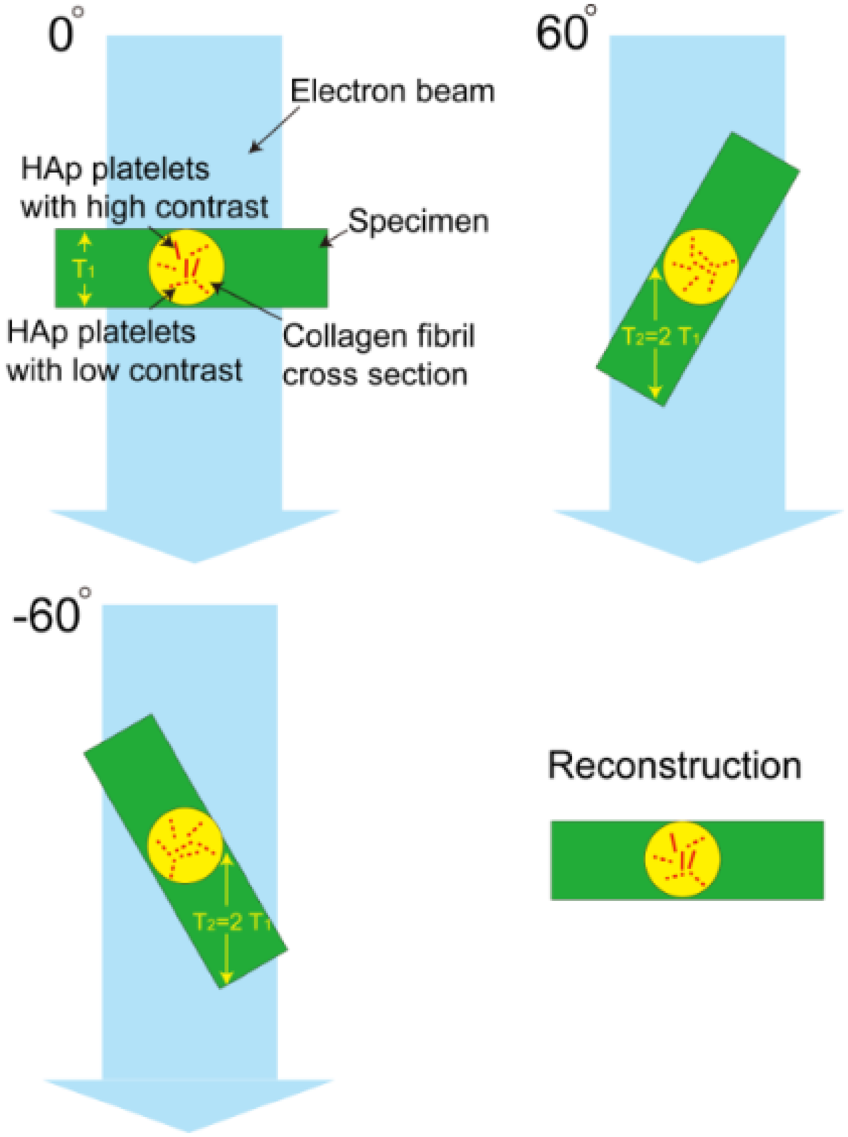


**Figure S6.** Contrast of HAp platelets in collagen fibrils at different tilting angles. At low tilting angle, e.g., 0°, the edge-on platelets are more visible, while the platelets in other orientations are low in contrast. At high tilting degrees, e.g., ±60°, however, all the platelets are lower in contrast as the actual sample thickness $T_2$ has doubled comparing with the thickness at 0°. As a result, the platelets that are edge-on at low tilting angles are more visible in the final reconstruction.

We can emulate this effect in our cryo-electron tomography study of the *in vitro* mineralized collagen fibril. For the same sample, in the area where ice layer is relatively thin (Fig. S7a, also Figures 2i to 2p in the main text), edge-on HAp platelets are observed at tilt angles of both 0° and 60°, although the image contrast is lower at 60°. In this case, indeed a random lateral orientation was observed for these platelets in the final reconstruction. In a thicker area (Fig. S7b), however, the image contrast at a tilt angle of 60° is very low. And in the reconstruction, indeed a pseudo "lateral orientation" was observed for the HAp platelets, with the angular distribution centered at 0°.

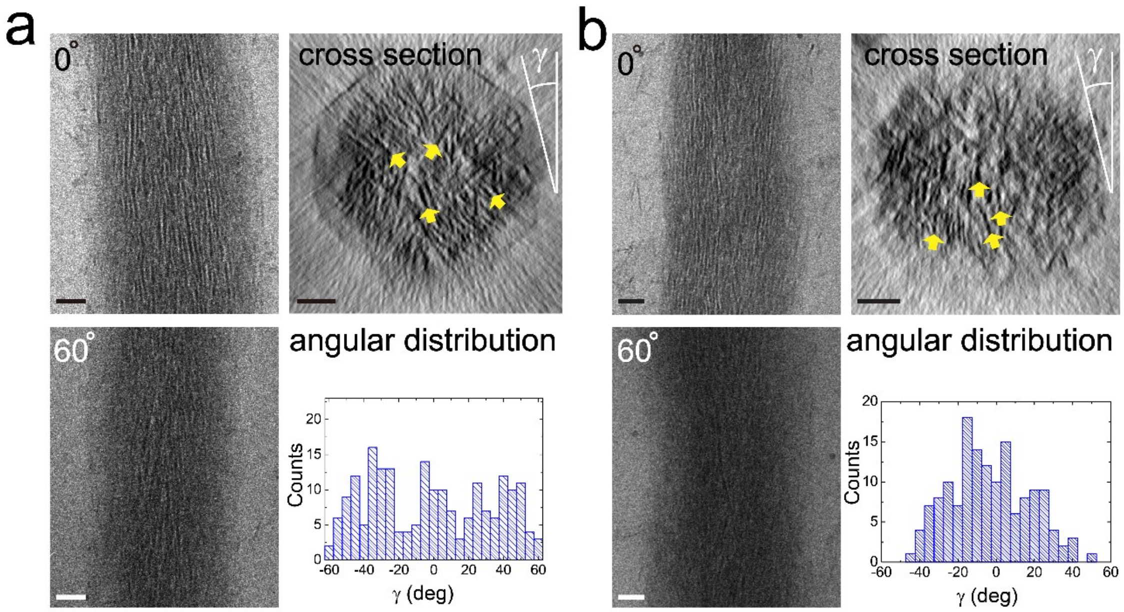


**Figure S7.** Sample thickness effect for cryo-electron tomography observation of collagen fibrils mineralized by HAp *in vitro*. (a) In a thinner area, edge-on HAp platelets are visible at both 0° and 60° tilting angles. In this case, no lateral orientation was observed for the platelets in the final reconstruction. (b) In a thicker area, edge-on platelets are still visible when tilting angle=0°. At 60°, however, the overall image contrast is very low, with the platelets hardly visible. As a result, in the final reconstruction, the platelets show a "lateral orientation", with the angular distribution center at 0°. Lateral orientation of several HAp platelets are highlighted by yellow arrows Scale bars: 50 nm.

In the 1993/1996 work of Landis et al. on tendon/chick bone,[1,2] the samples were ~500/~250 nm thick sections, which are both significantly thicker the sample used in our study (~100 nm thick). The TEM used for the previous two studies was a 1.0 MV Albany AEI-EM7 high voltage electron microscope, and the images were recorded using an EICONIX EC 78/99 digital camera with a pixel size of 1.3 nm. Although that the Albany AEI-EM7 has a higher accelerating voltage, the resolution/contrast achieved is unlikely to compete with the more recent FEI-Titan 300 kV TEM with large pole gap and energy filter that was used in our study, as the resolution in TEM is determined by many factors including the quality of electron source and lens, stability of stage, sensitivity of camera, use of energy filter, etc. Especially, at higher acceleration voltages the electron scattering cross-section of the specimen will be lower, which is actually not favored for beam-sensitive specimen with light elements (e.g., bone).[5] Indeed, in the 1996 work of Landis et al.,[1] the thickness of HAp platelets in bone was observed to be as high as 8 nm, suggesting a lack of resolution as a 2~4 nm thickness was observed in most of the recent studies.

The above shows that the intrafibrillar HAp platelets in bone are only uniaxially oriented, and explains that in the mid 1990's the state of the technology was not such that it could be observed in the pioneering experiments of Landis et al.

#### Supporting Information 7

##### Collagen sponge fibrils mineralized by HAp

Type-I collagen fibrils separated from collagen sponge (bovine Achilles) by grinding in liquid nitrogen and re-disperse in HEPES buffer were visualized by cryoTEM (Figure S8a), which shows ~300 nm thick fibrils with a clear 67 nm banding pattern, identical to what has been observed for self-assembled type I collagen fibrils (Horse Tendon).[6] The fibrils were mineralized by being exposed to a reaction solution containing $CaCl_2$, $K_2HPO_4$ and poly(aspartic acid) (pAsp). Significant intrafibrilar HAp mineralization was observed after 4 days (Figure S8b). LDSAED (inset of Figure S8b) shows a pair of narrow arcs corresponding to HAp (002) planes in the direction of collagen fibrils, together with 3 pairs of arcs with similar d-values corresponding to the (112), (221) and (300) diffractions, respectively. This observations is same with HAp mineralized horse tendon collagen (Figures 2i and 2l), and in-line with a previous report.[7]

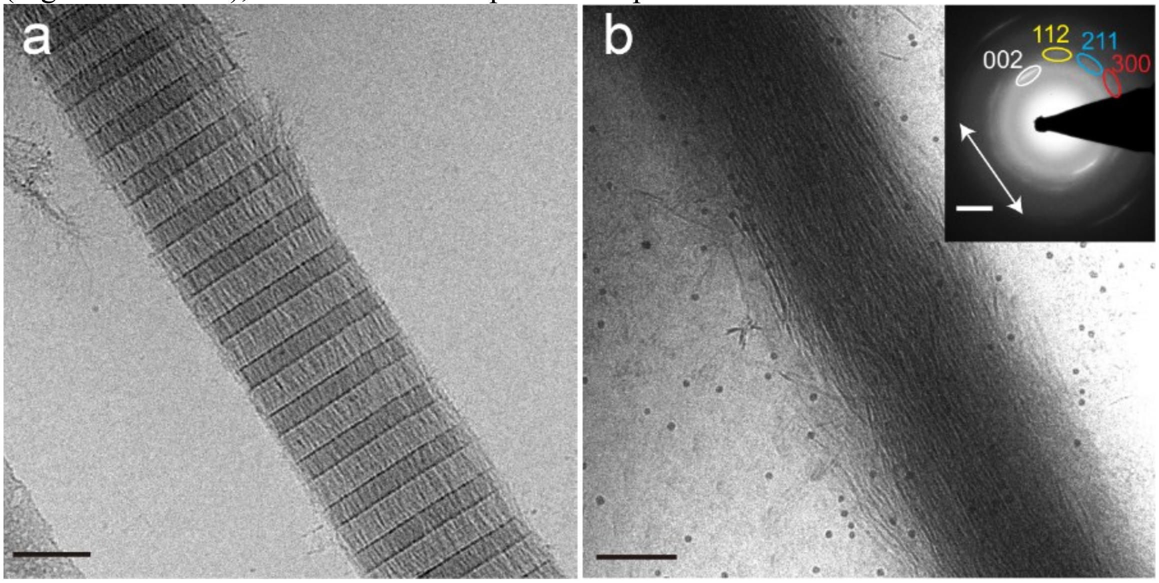


**Figure S8.** CryoTEM image of collagen fibrils (bovine Achilles) (a) before and (b) after HAp mineralization. Inset of (b): LDSAED pattern, with diffractions corresponding to HAp highlighted. The arrow indicates the direction of the collagen fibril. Scale bars: (a,b) 100 nm. Inset of (b): 2 $nm^{-1}$.

#### Supporting Information 8

##### Size of HAp crystals formed outside collagen fibril

We measured the size of HAp crystals that formed outside collagen fibril. Figure S9 shows a typical cryoTEM image, which was used to measure the size of the crystals. Only crystals found in the solution were measured, with length of ~130 nm, width of ~30 nm, thickness of ~4.5 nm and aspect ratio of ~4.3.

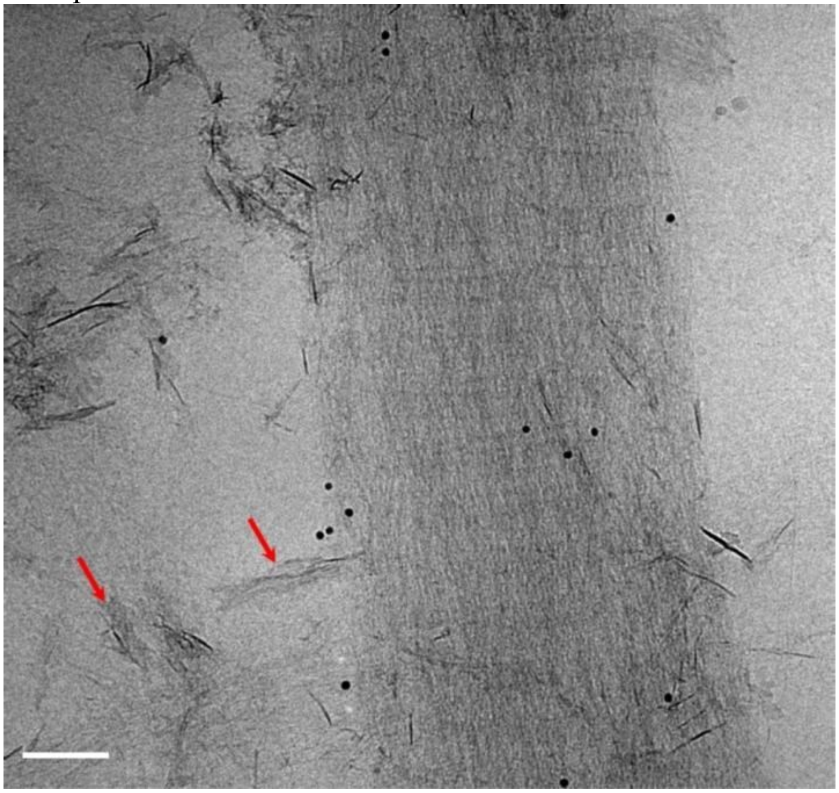

**Figure S9.** CryoTEM image of HAp crystals formed in solution. Arrows indicate representative crystals that were measured. Scale bar: 100 nm

#### Supporting Information 9

**pH profile of the diffusion of $NH_4(OH)$ into $Fe^{3+}/Fe^{2+}$ solution containing pAsp**

Collagen sponges were placed in $Fe^{3+}/Fe^{2+}$ solution containing pAsp (Fe/Asp = 0.25) and incubated overnight in a glove-box under $N_2$, saturated with 8% $NH_4(OH)$. Figure S10 shows the increase in pH over time, as a result of the diffusion of $NH_4(OH)$ into the solution.

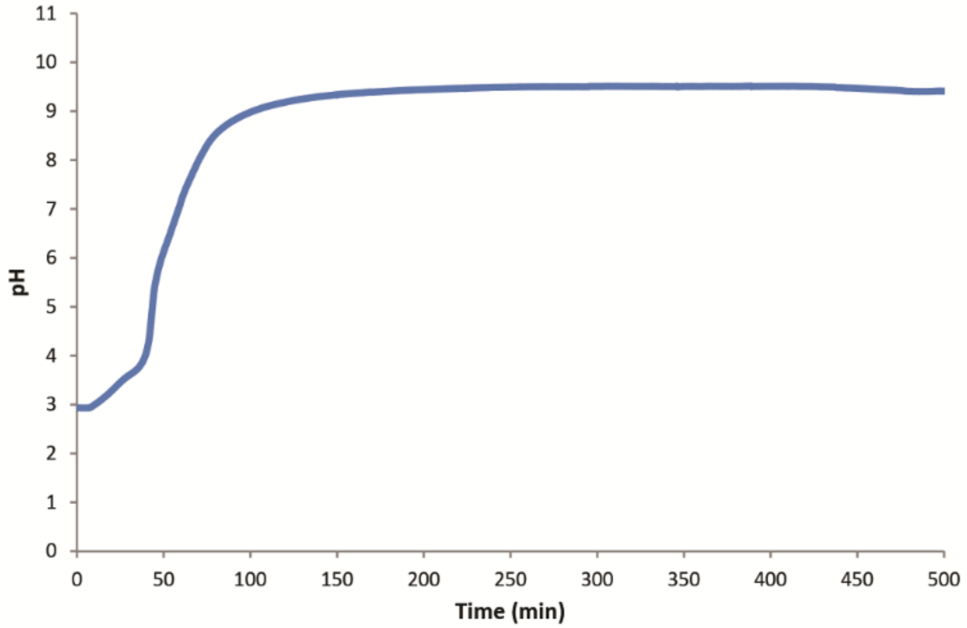


**Figure S10.** Increase of pH over time due to the diffusion of $NH_4(OH)$ 8% into a solution of $Fe^{3+}/Fe^{2+}$ containing pAsp (Fe/Asp = 0.25).

#### Supporting Information 10

##### Size of lepidocrocite crystals formed outside collagen fibril

We measured the size of lepidocrocite crystals that formed outside collagen fibril with a Fe: Asp ratio of 1:4. Figure S11 shows a typical dry TEM image, which was used to measure the size of the crystals. Only crystals outside the fibril were measured, with length of ~77 nm, width of ~25 nm, thickness of ~3 nm and aspect ratio of ~3.1

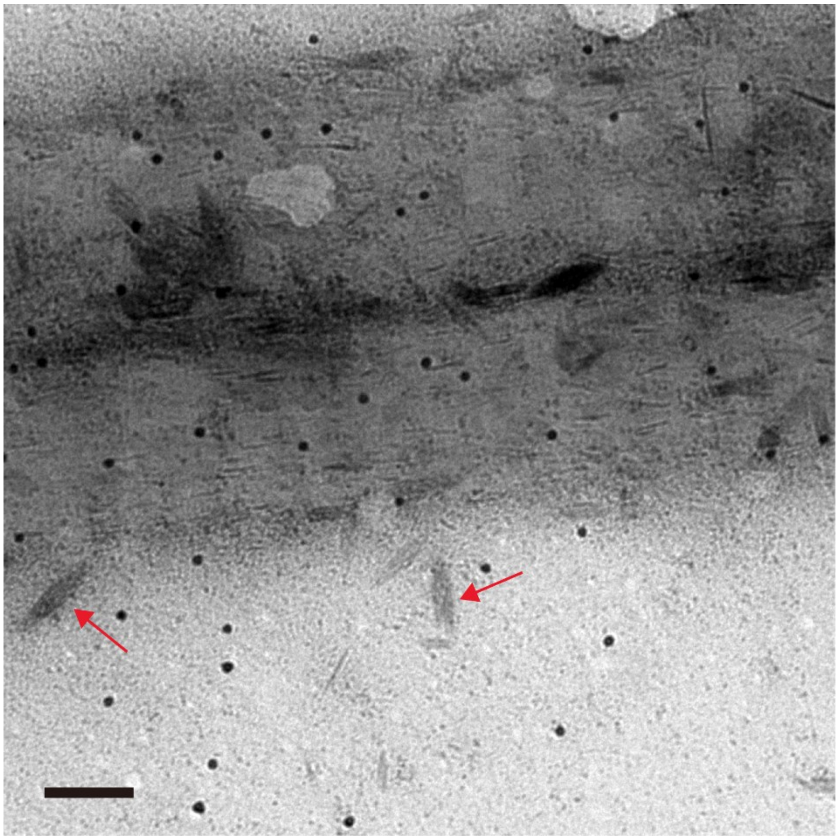

**Figure S11.** Dry TEM image of lepidocrocite crystals formed outside collagen fibril. Arrows indicate representative crystals that were measured. Scale bar: 100 nm.

#### Supporting Information 11

##### Iron oxide formation at different concentrations of pAsp

When the mineralization was performed without any additives, we obtained collagen fibrils that were completely coated with ferrihydrite particles 5-10 nm in size (Figures S12a and S12b). With a ratio of Fe: Asp of 4:1, the collagen was still covered with ferrihydrite, however few needle-shaped crystals were visible, with their long axis aligned in the direction of the long axis of the fibril (Figures S12c and S12d). When the concentration of pAsp was increased to a ratio of 1:1 Fe: Asp, ferrihydrite did not form in the reaction anymore. Instead, lepidocrocite was present, both randomly oriented and with their long axis aligned in the direction of the collagen fibril (Figures S12e and S12f). By increasing the concentration of pAsp to a ratio of 1:4 Fe: Asp, only oriented lepidocrocite crystals were visible (Figures S12g and S12h).

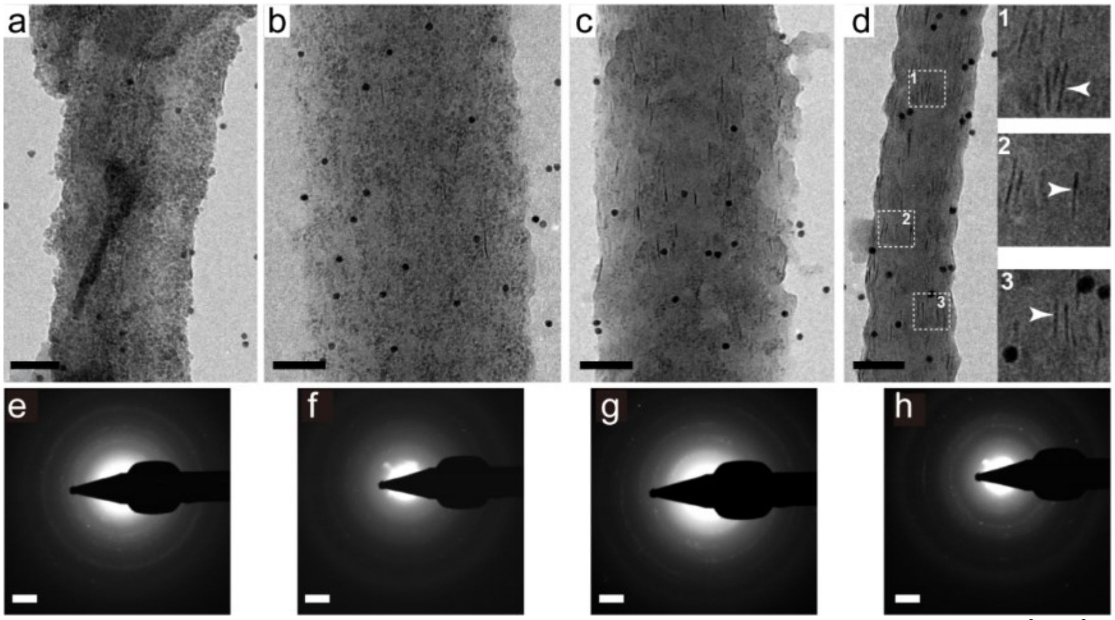


**Figure S12.** (a, c, e, g). Dry TEM image of collagen fibril mineralized by co-precipitation of $Fe^{3+}/Fe^{2+}$ ions, with Fe:Asp ratio of 1:0, 4:1, 1:1 and 1:4, respectively. Insets of (g): Zoom-in images showing the lepidocrocite crystals aligned along the long axis of collagen (arrowheads). (b, d, f, h) LDSAED of (a, c, e, g), respectively. (b) and (d) correspond to 2-line ferrihydrite, while (f) and (h) shows the formation of lepidocrocite. Scale bars: (a, c, e, g) 100 nm. (b, d, f, h) 2 $nm^{-1}$.

#### Supporting Information 12

##### Low dose selected area electron diffraction analysis of collagen sponge mineralized with lepidocrocite in the presence of pAsp

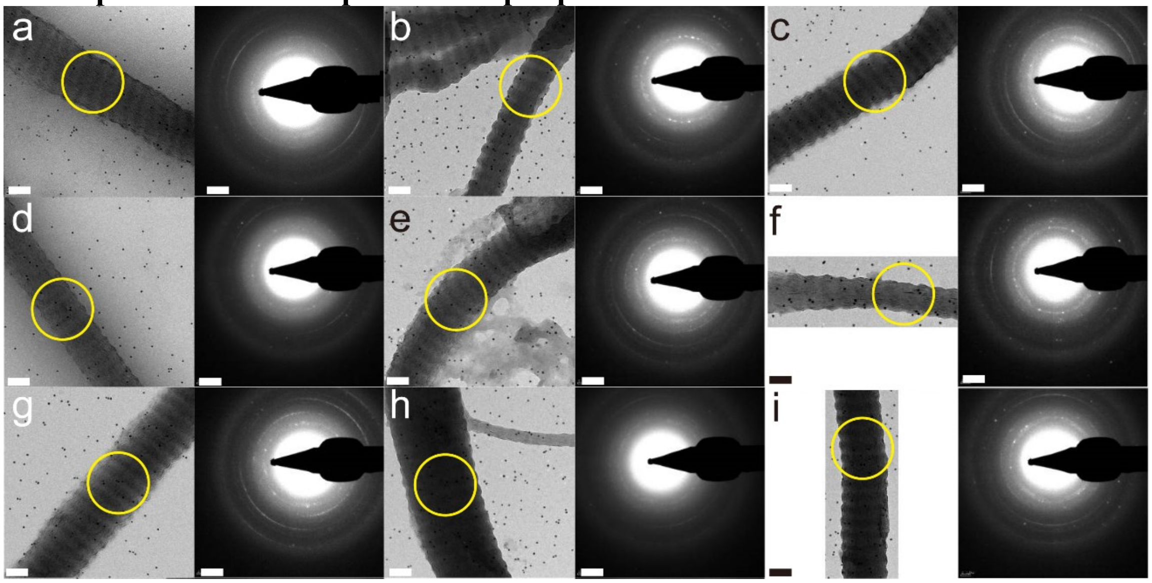


**Figure S13.** (a-i) Summary of 9 TEM images and corresponding LDSAED patterns of collagen mineralized with lepidocrocite in the presence of polyaspartic acid. The areas where the LDSAED patterns were taken are highlighted by yellow circles. Scale bars: TEM images: 100 nm. LDSAED patterns: 2 $nm^{-1}$.

The 9 LDSAED patterns (Figure S13) were radially averaged. These 9 averages were again averaged to give a 1D diffraction spectrum as shown in Figure S14. The peak positions and corresponding reflections are summarized in Table S1.The reflections were assigned to lepidocrocite with the help of the XRD data reported by Ewing.[8]

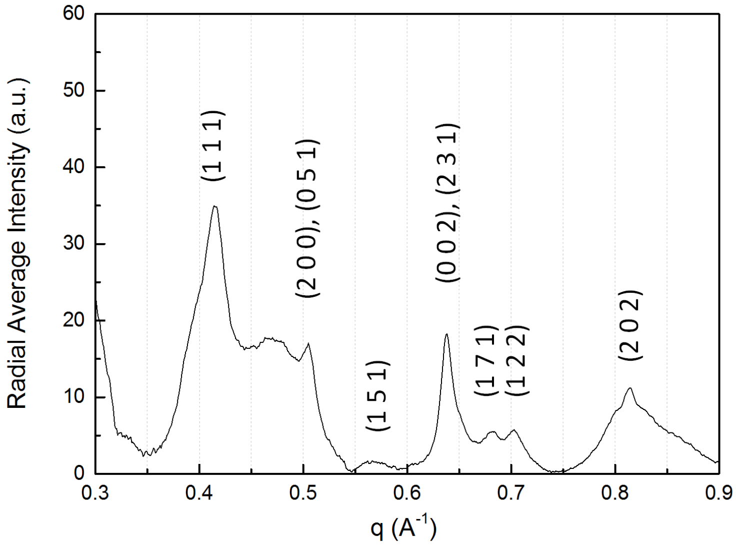


**Figure S14.** Radially averaged diffraction data from the average of all LDSAED patterns.

**Table S1.** Summary of the positions of diffraction peak maxima and corresponding reflections.

| Reciprocal Distance ($Å^{-1}$) | Distance(Å) | Reflection |
|---|---|---|
| 0.423 | 2.366 | (1 1 1) |
| 0.480 | 2.09 | (0 6 0), (1 3 1) |
| 0.516 | 1.94 | (0 5 1) |
| 0.517 | 1.93 | **(2 0 0)** |
| 0.578 | 1.729 | (1 5 1) |
| 0.652 | 1.534 | **(0 0 2)**, (2 3 1) |
| 0.696 | 1.436 | (1 7 1), (1 8 0) |
| 0.718 | 1.394 | (1 2 2) |
| 0.832 | 1.203 | (2 0 2) |

To generate the averaged diffraction pattern used in Figure 2c, the center of the individual diffraction patterns (Figure S13) was determined by fitting a ring on the (002) wedge and the orientation was deduced from the corresponding TEM images. Subsequently, these diffraction patterns were translated and rotated with respect to their center and orientation resulting in aligned diffraction patterns. From these, the beamstop was removed by fitting a mask and the diffraction patterns were normalized with the average intensity in the 0.3-0.9 $Å^{-1}$ range. Next, the diffraction pattern was calculated and the background was subtracted with the *imtophat* function in Matlab using a disk of ~ 0.1 $Å^{-1}$ as structuring element. This yielded the final averaged diffraction pattern, with the all reflections aligned to give the collagen a vertical orientation, with respect to the averaged diffraction pattern.

#### Supporting Information 13

**Collagen sponges mineralized by $CaCO_3$ without the presence of polymeric additives**
When the collagen sponges were mineralized without polymeric additives, rhombohedral calcite crystals were found between the collagen fibrils (Figure S15a). Those crystals show a higher brightness in the back-scattered electron image comparing to the fibrils, indicating that no $CaCO_3$ was formed within the fibrils (Figure S15b). This was confirmed by energy dispersive X-ray spectroscopy (EDX, Figure S15c), which shows no Ca signal in the collagen fibrils.

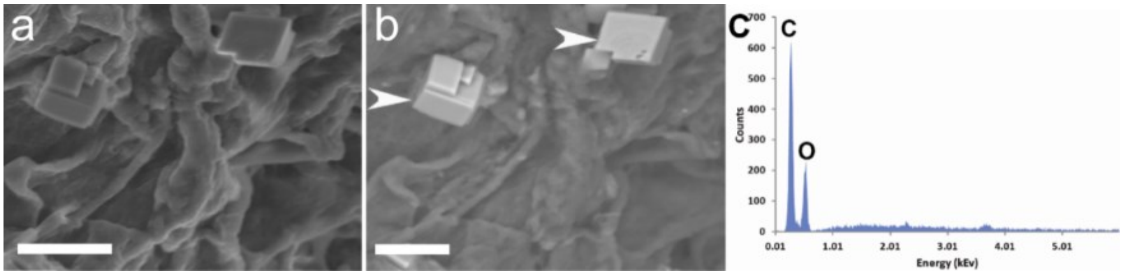


**Figure S15.** SEM images of the collagen sponge mineralized with $CaCO_3$ without the presence of polymeric additives. (a) Secondary electrons detector image of the collagen sponge. (b) Back-scattered electrons detector image of (a) White arrowheads: calcite crystals found between the collagen fibrils. (c) EDX analysis of (a), showing the absence of Ca in the collagen fibrils. Scale bars: 2 μm.

#### Supporting Information 14

**Experimental setup for the time-resolved measurements of the mineralization of collagen sponges with $CaCO_3$ using SAXS and WAXS**

To perform *in situ* SAXS and WAXS study on the mineralization of collagen sponges with $CaCO_3$, 0.125 $cm^3$ of collagen sponge (bovine Achilles) was incubated in a flow cell between two mica windows. Subsequently, a solution of 10 mM $CaCl_2$ containing 1 mg $mL^{-1}$ of pAH in a desiccator containing $(NH_4)_2CO_3$ was pumped through the flow cell with an addition rate of 2.5 ml/min. The flow cell was analyzed using SAXS/WAXS. The data were acquired every 5 minutes with an acquisition time of 3 minutes, over an 8 h period. The X-ray wavelength was set to $\lambda = 0.1$ nm and the sample-to-detector distances were 130 cm and 28 cm for SAXS and WAXS, respectively. The q-range (where $q = 4\pi/\lambda \sin(\theta)$ is the modulus of the scattering vector, and $\theta$ is half of the scattering angle) was calibrated according to the position of diffraction peaks from Silver Behenate and $\alpha$-Alumina standard samples. High sensitivity, noiseless photon counting Pilatus detectors (Pilatus1M for SAXS and Pilatus 300K for WAXS) were used to collect 2D images. Standard corrections for beam intensity, empty cell background and sample transmission have been applied before 2D images integration.

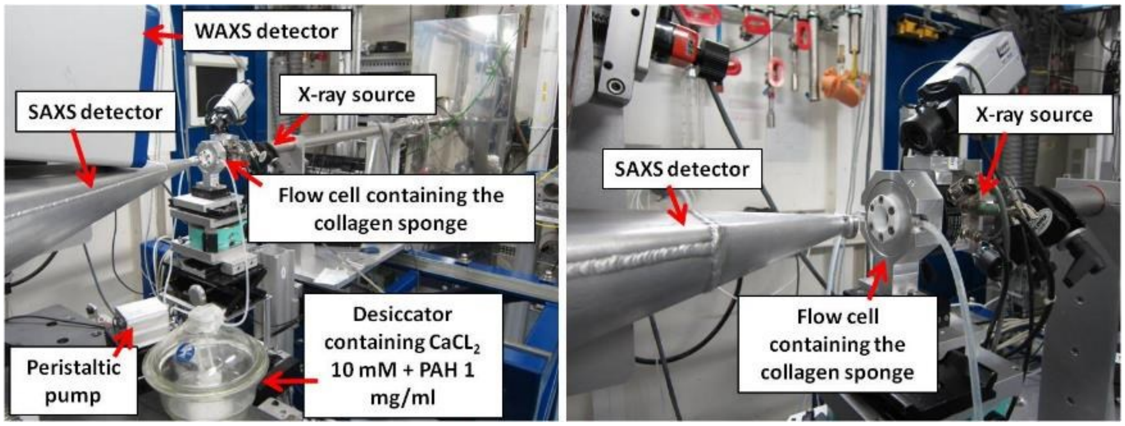


**Figure S16.** Experimental setup for measuring simultaneously SAXS and WAXS spectra during the mineralization of collagen sponges with $CaCO_3$ in the presence of pAH.

#### Supporting Information 15

**Influential books/papers presenting the 'deck-of-cards' model**

Some of the influential books/papers presenting the 'deck-of-cards' model of mineralized collagen fibril in bone are listed below together with the citation numbers. All the citation numbers are based on Google Scholar records on 20, Mar. 2020:

1. Lowenstam, H. A. and S. Weiner (1989). On biomineralization, Oxford University Press on Demand. (Cited: 3486)
2. Weiner, S. and H. D. Wagner (1998). "The material bone: Structure mechanical function relations." Annual Review of Materials Science **28**: 271-298. (Cited: 2660)
3. Rho, J.-Y., et al. (1998). "Mechanical properties and the hierarchical structure of bone." **20**(2): 92-102. (Cited: 2412)
4. Mann, S. (2001). Biomineralization: principles and concepts in bioinorganic materials chemistry, Oxford University Press on Demand. (Cited: 2709)
5. Dorozhkin, S. V. and M. J. A. C. I. E. Epple (2002). "Biological and medical significance of calcium phosphates." **41**(17): 3130-3146. (Cited: 1838)
6. Gao, H. J., et al. (2003). "Materials become insensitive to flaws at nanoscale: Lessons from nature." Proceedings of the National Academy of Sciences of the United States of America **100**(10): 5597-5600. (Cited: 1569)
7. Currey, J. D. (2006). Bones: structure and mechanics, Princeton university press. (Cited: 1957)
8. Fratzl, P. and R. Weinkamer (2007). "Nature's hierarchical materials." Progress in materials Science **52**(8): 1263-1334. (Cited: 1897)
9. Meyers, M. A., et al. (2008). "Biological materials: structure and mechanical properties." **53**(1): 1-206. (Cited: 1874)
10. Wegst, U. G., et al. (2015). "Bioinspired structural materials." **14**(1): 23-36. (Cited 1672)

**Movie S1.** Electron tomographic reconstruction y-slices of the human bone lamella, showing the overview of the sample.

**Movie S2.** Electron tomographic reconstruction y-slices of the two mineralized collagen fibrils in human bone as shown in Figures 2a to 2h.

**Movie S3.** Electron tomographic reconstruction z-slices of the two mineralized collagen fibrils in human bone as shown in Figures 2a to 2h.

**Movie S4.** 3D electron density map of collagen structure created from X-ray diffraction data. The white fibrils correspond to the collagen molecules, and the channels in the gap region are labelled by different colors based on their connectivity.

**Movie S5.** CryoEM tomographic reconstruction y-slices of a collagen fibril mineralized *in vitro* by HAp as shown in Figures 2i to 2p.

**Movie S6.** CryoEM tomographic reconstruction z-slices of a collagen fibril mineralized *in vitro* by HAp as shown in Figures 2i to 2p.

**Movie S7.** EM tomographic reconstruction y-slices of a collagen fibril mineralized *in vitro* by lepidocrocite as shown in Figure 4.

**Movie S8.** EM tomographic reconstruction z-slices of a collagen fibril mineralized *in vitro* by lepidocrocite as shown in Figure 4.

#### References


1 Landis, W. J., Hodgens, K. J., Arena, J., Song, M. J. & McEwen, B. F. Structural relations between collagen and mineral in bone as determined by high voltage electron microscopic tomography. *Microsc. Res. Tech.* **33**, 192-202 (1996).

2 Landis, W. J., Song, M. J., Leith, A., Mcewen, L. & Mcewen, B. F. Mineral and Organic Matrix Interaction in Normally Calcifying Tendon Visualized in 3 Dimensions by High-Voltage Electron-Microscopic Tomography and Graphic Image-Reconstruction. *J. Struct. Biol.* **110**, 39-54 (1993).

3 Friedrich, H., de Jongh, P. E., Verkleij, A. J. & de Jong, K. P. Electron tomography for heterogeneous catalysts and related nanostructured materials. *Chem. Rev.* **109**, 1613-1629 (2009).

4 Reznikov, N., Bilton, M., Lari, L., Stevens, M. M. & Kröger, R. Fractal-like hierarchical organization of bone begins at the nanoscale. *Science* **360**, eaao2189 (2018).

5 Cosslett, V. E. High voltage electron microscopy and its application in biology. *Philosophical Transactions of the Royal Society of London. B, Biological Sciences* **261**, 35-44 (1971).

6 Nudelman, F. *et al.* The role of collagen in bone apatite formation in the presence of hydroxyapatite nucleation inhibitors. *Nat. Mater.* **9**, 1004-1009 (2010).

7 Olszta, M. J. *et al.* Bone structure and formation: A new perspective. *Mater. Sci. Eng. R.* **58**, 77-116, doi:10.1016/j.mser.2007.05.001 (2007).

8 Ewing, F. The crystal structure of lepidocrocite. *J. Chem. Phys.* **3**, 420-424 (1935).
